# Improved AlphaFold modeling with implicit experimental information

**DOI:** 10.1101/2022.01.07.475350

**Authors:** Thomas C. Terwilliger, Billy K. Poon, Pavel V. Afonine, Christopher J. Schlicksup, Tristan I. Croll, Claudia Millán, Jane. S. Richardson, Randy J. Read, Paul D. Adams

## Abstract

Machine learning prediction algorithms such as AlphaFold^1^ and RoseTTAFold^2^ can create remarkably accurate protein models, but these models usually have some regions that are predicted with low confidence or poor accuracy^3–6^. We hypothesized that by implicitly including experimental information, a greater portion of a model could be predicted accurately, and that this might synergistically improve parts of the model that were not fully addressed by either machine learning or experiment alone. An iterative procedure was developed in which AlphaFold models are automatically rebuilt based on experimental density maps and the rebuilt models are used as templates in new AlphaFold predictions. We find that including experimental information improves prediction beyond the improvement obtained with simple rebuilding guided by the experimental data. This procedure for AlphaFold modeling with density has been incorporated into an automated procedure for crystallographic and electron cryo-microscopy map interpretation.

---

Advanced machine learning-based structure prediction algorithms are transforming the way that three-dimensional structures of proteins and their complexes are obtained^3,4,7,8^. The AlphaFold^1^ and RoseTTAFold^2^ algorithms, for example, can often create accurate models for substantial regions of a protein structure based on the amino acid sequence of that protein and on residue covariation information^9^ present in a multiple sequence alignment^1^. Prediction can be augmented by including experimentally determined structures of proteins with similar sequences as templates^1^. In many cases the resulting predicted models are accurate enough to allow straightforward structure determination using molecular replacement in macromolecular crystallography or by docking a structure in a density map in single-particle cryo-electron microscopy (cryo-EM), without requiring that a similar structure has been previously determined^7,8,10^.

There are limitations in using predicted models for structure determination^3–5^. In particular, machine-learning methods typically do not yield accurate predictions for all of the residues in a protein^6^. This is partly due to the presence of disordered segments in many proteins^1,11^, but is also due to the limited size and accuracy of multiple sequence alignments for part or all of some protein sequences, resulting in a limited amount of available information about residue covariation ^1^. A related limitation is that parts of proteins that can adopt alternative conformations may be systematically predicted in only one of them^1,12^; this limitation may be reduced by alternative sampling of multiple sequence alignments^12^. Additionally, individual domains of proteins are often predicted accurately, but in the absence of extensive conserved interaction surfaces the spatial relationship between domains cannot be unambiguously predicted with current methods^3^. A final limitation is that as these machine learning methods are trained on structures in the PDB^1^, predictions are likely to be biased towards these known structures even if they are not included explicitly as templates in prediction.

A strength of recent machine-learning algorithms for protein structure prediction is that they can assess the accuracies of their own predictions. AlphaFold, for example, estimates the value of a commonly-used measure of model accuracy (lDDT-C_α_^13^) for each residue in a protein and reports these estimates as a confidence measure, plDDT^1^. Validation with known structures demonstrated that these AlphaFold plDDT values are reasonably good indicators of actual accuracy (Pearson’s r value relating plDDT and lDDT-C_α_ is 0.73^6^).

It is well known that the accuracy of structure prediction can be improved by including external structural information, for example distances between specified pairs of residues in a protein^14^. In AlphaFold and RoseTTAFold, for example, residue pair distance information is implicitly derived from sequence covariation^1,2^. It is reasonable to expect that experimental structural information from density maps such as those used in cryo-EM or crystallographic structure determination could be included as well, though a mechanism for incorporation of this information in a form that is compatible with modeling would be required.

The hypothesis underlying the present work is that experimental information might improve structure prediction synergistically, where correcting one part of a protein chain might improve structure prediction in another part of the chain. In AlphaFold, a core algorithm focuses attention on features that may contribute the most to structure prediction^1^. An internal recycling procedure uses the path of the protein chain in one cycle to focus attention on interactions that should be considered in the next cycle. If experimental information were to result in adjustments in conformation, the attention mechanism might recognize important relationships that otherwise would have been missed. This means that experimental information might be amplified by the prediction algorithm. At the same time, improvement in the accuracy of a predicted model might make it easier to identify modifications to that model needed to obtain a better match to the density map. These possibilities suggest that an iterative procedure for incorporation of information from a density map into structure prediction might further improve the accuracy of modeling. This would be similar to the situation in macromolecular crystallography, where improvement of one part of a model leads to improved estimates of crystallographic phases, in turn improving the density map everywhere and allowing still more of the model to be built^15^.

A second hypothesis in this work is that information from a density map can be partially captured in the form of a rebuilt version of a predicted model that has been adjusted to match the map. The structure of such a model could only represent a small part of the total information in a map, but it seemed possible that much of the key information could be captured, including overall relationships between domains in a protein as well as the detailed conformation of the protein. As AlphaFold can use models of known proteins as templates^1^, such a rebuilt model could readily be incorporated into subsequent cycles of structure prediction.

We tested these ideas by developing an automated procedure in which a predicted AlphaFold model is trimmed, superimposed (docked) on a cryo-EM density map, and rebuilt to better match the map. The rebuilt model is then supplied along with the sequence to AlphaFold in a new cycle of prediction. The output of this procedure is a new AlphaFold model that has incorporated experimental information through the use of the rebuilt template in the prediction. We applied four cycles of the iterative algorithm to the sequence of one protein chain and the full density map for each of 25 cryo-EM structures, all deposited after the training database for the version of AlphaFold we used was created (July 2020). In these tests, multiple sequence alignments were included in each stage of AlphaFold modeling. To emulate the situation where no similar structure is present in the PDB, templates from the PDB were not used. For each protein we then examined the four AlphaFold models obtained (one for each cycle of modeling), comparing them to the corresponding deposited model (used as our best estimate of the true structure) and to the corresponding deposited density map.

Fig. 1 (A-F) illustrates iterative structure prediction for one of these structures, that of a focused reconstruction of SARS-CoV-2 spike protein receptor binding domain (RBD) in a complex with neutralizing antibodies^16^ (3.7 Å; EMDB entry 23914, PDB entry 7mlz; only the spike protein is analyzed here). Panel A shows that the five-stranded ß-sheet (lower left corner) in the AlphaFold model (in blue) created based on the sequence of the spike protein can be superimposed closely on the deposited model (in brown), but the loops near P479 in the right part of (A) then do not match well. The same AlphaFold model is shown along with the density map in (**D**), where it can be appreciated that the density map does not clearly show the path of the protein chain. The agreement between the AlphaFold model and the map is considerably worse than between the deposited model and map (map correlation with map calculated from deposited model is 0.70; from AlphaFold model is 0.41). Panel B shows a rebuilt version of this AlphaFold model (in purple) obtained after automatic rebuilding using the density map. It is different from the blue predicted model in panel A and agrees better with the density map (panel E), where the map correlation increased from 0.41 to 0.58. The percentage of C_α_ atoms in the deposited model matched within 3 Å by a C_α_ atom in the rebuilt model was also somewhat improved over that for the superimposed AlphaFold model (from 71% to 76%). This rebuilt model was used as a template in AlphaFold modeling, with the goal of providing the inference procedure with some additional information about which parts of the structure are close together, and the rebuilding and modeling were repeated for a total of four iterations. The AlphaFold model obtained after iterative prediction and rebuilding is shown in green in panel C. It matches the deposited model (in brown) much more closely than the original AlphaFold model obtained with sequence alone, particularly in the loop region near residue P479, and 91% of C_α_ atoms in the deposited model were matched within 3 Å by a C_α_ atom in the superimposed AlphaFold model. The overall map correlation for AlphaFold model obtained after iterative prediction and rebuilding is 0.57. Note that unlike the rebuilt model, the AlphaFold predicted model shown in panel C has not been adjusted by coordinate refinement or rebuilding; it is simply superimposed as a rigid unit on the density map. The similarity obtained to the map and to the deposited model therefore reflects an improvement in the AlphaFold prediction itself.

**Fig. 1.**
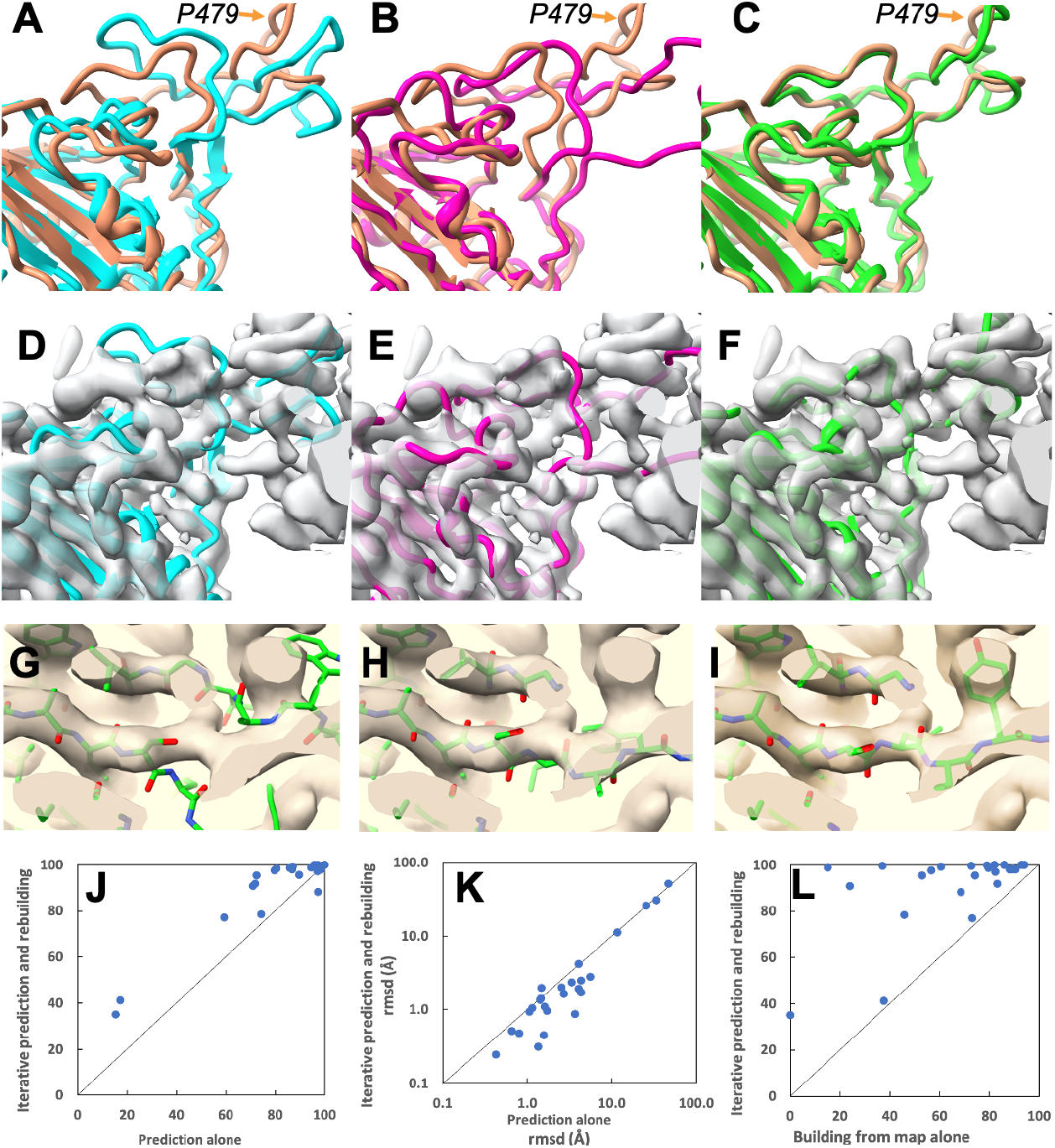
Iterative AlphaFold prediction and model rebuilding using density maps. **A**. Comparison of AlphaFold model of SARS-CoV-2 spike protein receptor binding domain (blue) with deposited model^16^ (PDB entry 7mlz, brown). The position of V445 is indicated. B. Comparison of model in A rebuilt using density map (in purple) with deposited model (brown). C. AlphaFold model obtained using density map and four cycles of iteration including rebuilt models as templates (green), compared with deposited model (brown). D, E, F. Models as in A, B and C, superimposed on the map used for rebuilding (EMDB entry 23914^17^, automatically sharpened as described in Materials and Methods). G, H, I. Details of iterative rebuilding of the 2AG3 Fab heavy chain^18^ (PDB entry 7mjs chain H) using cryo-EM data from EMDB entry 23883 at a resolution of 3.0 Å. G. AlphaFold prediction superimposed on density map. H. AlphaFold prediction as in G, but after one cycle of iterative rebuilding. I, As in H, but after 4 cycles of iterative rebuilding. J. Accuracy of models obtained with AlphaFold alone (abscissa) and obtained with iterative AlphaFold prediction and rebuilding with density (ordinate) for one chain from each of 25 structures from the PDB and EMDB. Accuracy is assessed as the percentage of of C_α_ atoms in the deposited model matched within 3 Å by a C_α_ atom in the superimposed AlphaFold model. K. Accuracy of models shown in J, assessed based on rmsd of matching C_α_ atoms and shown on a log scale. Abscissa is rmsd for models obtained with AlphaFold alone and ordinate is for models obtained with iterative AlphaFold prediction and rebuilding with density. L. Accuracy of models assessed as in J by the percentage of of C_α_ atoms in the deposited model matched within 3 Å by a C_α_ atom in the superimposed model, obtained with direct model-building using the corresponding density maps using the Phenix tool map_to_model (abscissa) compared with those obtained with iterative AlphaFold prediction (ordinate).

Overall, panels A-F of Fig. 1 show that the AlphaFold model obtained with our iterative procedure and shown in green in panel C is much more similar to the deposited model (brown) than is either the predicted AlphaFold model created with sequence alone, shown in blue in panel A, or the rebuilt version of this predicted model, shown in purple in panel B. The improvement over the original AlphaFold model supports the idea that a template created by rebuilding an AlphaFold model using a density map contains information from that density map that can be used to improve AlphaFold structure prediction. The observations that the AlphaFold model obtained using a density map also improves upon the rebuilt model and that iteration improves the AlphaFold model support the idea that model rebuilding is synergistic with AlphaFold prediction, yielding a new model that is better than either alone.

Panels G, H and I of Fig. 1 illustrate the improvement of another AlphaFold prediction by iterative rebuilding and modeling. A detail of the superimposed AlphaFold prediction of the 2AG3 Fab heavy chain^18^ is shown in panel G along with the corresponding portion of the density map from EMDB entry 23883 at a resolution of 3.0 Å. The superimposed predicted model does not match the density well, and the rmsd of all matching C_α_ atoms from the deposited model is 3.6 Å. Panel H shows that the superposed AlphaFold model obtained after one cycle of iterative rebuilding matches the map considerably better, and panel I shows that after four cycles the AlphaFold model closely matches the density map. The full AlphaFold prediction for this heavy chain obtained after four cycles of iteration with the density map has an rmsd of matching C_α_ atoms from the deposited model of just 0.8 Å.

Panel J compares the accuracy of AlphaFold models obtained without and with density information for all of the 25 recently-deposited structures considered. The inclusion of density information increased the number of these 25 structures with at least 90% of C_α_ atoms superposing within 3 Å from 12 to 20. This set of models is assessed based on rmsd of matching C_α_ atoms in panel K demonstrating that in most cases the iterative AlphaFold models have much lower rmsd from corresponding deposited models than predictions using sequence alone.

Panel L of Fig. 1 extends this analysis further by comparing the 25 models obtained using iterative AlphaFold modeling and model rebuilding (abscissa) with models created directly from density maps using an automatic model-building algorithm^19^ that is based on many of the same tools used here in model rebuilding, but without including AlphaFold at all (ordinate). All but one of the iterative AlphaFold models are more accurate than the corresponding models created by automatic model-building alone.

In the cases described above, it was possible that information about the specific sequences that are being modeled could be present in the AlphaFold parameter database because similar structures may have been present in the PDB when AlphaFold was trained. In this work we are comparing AlphaFold predictions that are identical except that they are carried out with and without templates, so this does not directly affect our conclusion that AlphaFold modeling and rebuilding using a density map are synergistic. There was a possibility, however, that including the density information in these examples allowed AlphaFold prediction to use some pre-existing information about similar structures, rather than truly incorporating new information from the density maps. To address such a possibility, we carried out an analysis of a structure for which no similar structure was present in the PDB when AlphaFold training was carried out. The structure we used was that of a domain of a bacterial flagellar basal body^20^ (PDB entry 7bgl, chain a, residues 250-365, EMDB entry 12183, resolution of 2.2 Å) included in the CASP-14 structure prediction competition^21^ (target identification of T1047s2-D3). The PDB entry with the most similar sequence (PDB entry 2hm2) present at the time of AlphaFold training has a sequence identity of just 9% and has a very different structure^21^. Parts of the structure of this domain from the basal body are accurately predicted by AlphaFold^21^, however there was a substantial difference in the arrangement of two antiparallel strands relative to the cryo-EM structure, as well as a small difference in the position of a helix (cf. panel A of Fig. 2 and compare the four-stranded sheet in the AlphaFold model in blue with the two-stranded sheet in the deposited model in brown at the left side of the figure, and compare positions of the blue and brown helices in the center).

**Fig. 2.**
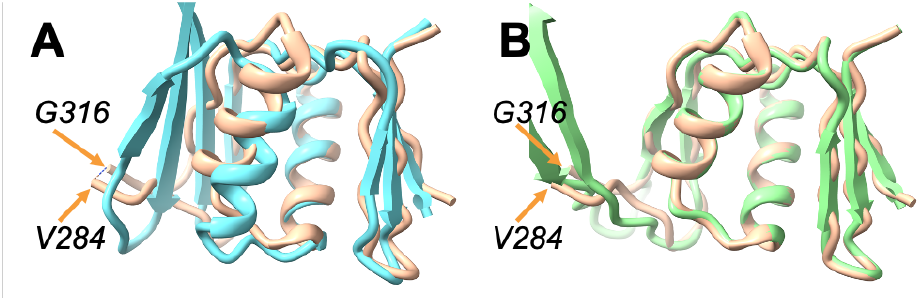
Iterative AlphaFold prediction and model rebuilding of domain from flagellar basal body. A. Comparison of AlphaFold model flagellar basal bodychain a residues 250-365 (blue) with deposited model^20^ (PDB entry 7bgl, brown). The positions of G316 and V284, bracketing a segment that is not present in the deposited model, are indicated. B. Comparison of model in A obtained with three cycles of iterative AlphaFold modeling and rebuilding using density map (in green) with deposited model (brown).

We used the flagellar basal body (7bgl) structure to test whether iterative AlphaFold prediction and model rebuilding would be effective in a case where AlphaFold was trained without any similar structures. In this test, fragments from a model automatically built from the density map were included in model rebuilding, and multiple sequence alignments were only used in the first cycle of AlphaFold modeling. These options were chosen to improve model rebuilding and to allow the conformations of the rebuilt models to guide the AlphaFold prediction. Panel A of Fig. 2 showed that a standard AlphaFold prediction leads to a model that has some correct and some substantially incorrect parts. Note that the deposited model in brown is missing residues 285-315 which are not visible in the density map. These residues are modeled by AlphaFold but are not included in our comparisons. Iteration of AlphaFold modeling with model rebuilding yields a model that agrees more closely with the deposited (7bgl) model (panel B) This iterative AlphaFold model is much more accurate than the original AlphaFold prediction (panel A) based on rmsd between matching C_α_ atoms (1.7 Å vs 4.7 Å) and by percentile-based spread, which deemphasizes large discrepancies^22^ (0.3 Å vs 2.0 Å). It is similar to but somewhat more accurate than the initial rebuilt model (rmsd of 1.8 Å, percentile-based spread of 0.4 Å). To check that the improvement in prediction with iteration was not simply due to leaving out the multiple sequence alignment in predictions after the first, we carried out AlphaFold modeling without a multiple sequence alignment and without information from the map. This resulted in a prediction that was quite different from that of the deposited model (rmsd of 11.5 Å, percentile-based spread of 11.2 Å). These observations show that the synergy in iterative AlphaFold modeling and model rebuilding using a density map can be obtained even if AlphaFold is trained in the absence of any similar structures.

An immediate application of iterative prediction and model rebuilding is automatic analyses of cryo-EM or crystallographic density maps. Though tools exist for this purpose, automatic map interpretation is challenging, particularly when high-resolution maps are not available. For example, Panels A-D of Fig. 3 show that automatically-generated models created by each of two automated tools ^19,23^ using the experimental density map for the SARS-CoV-2 spike protein structure illustrated in Fig. 1 fail to create a model resembling the deposited structure. The automated map interpretation methods in panels A and C are able to match just 60% and 24%, respectively, of C_α_ atoms in the corresponding deposited model within 3 Å.

**Fig. 3.**
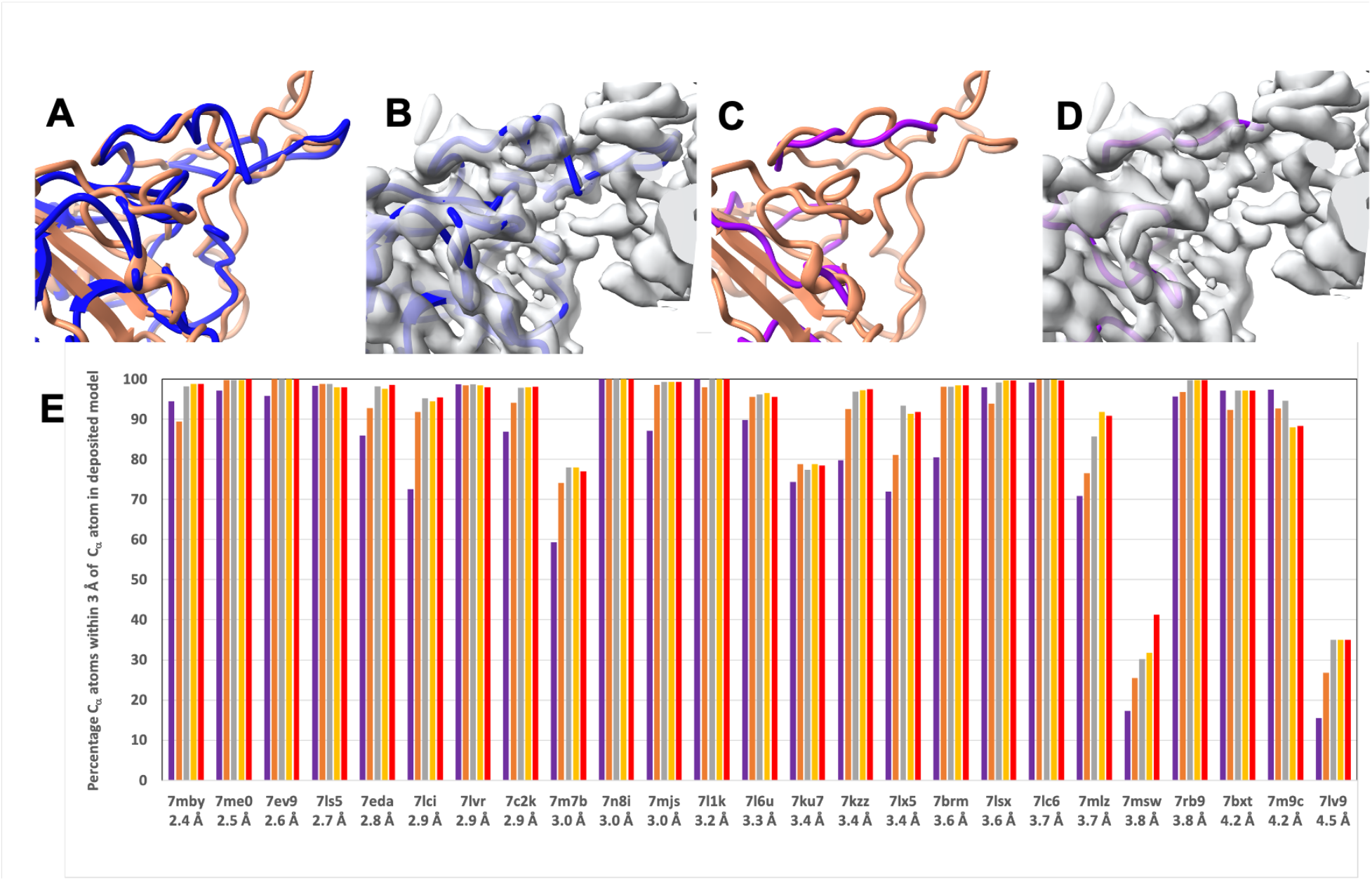
Current automatic map interpretation tools work poorly with an unclear map but can be improved upon by iterative AlphaFold prediction and model rebuilding. A. Machine-learning method for automatic map interpretation (DeepTracer^23^) applied to the SARS Cov-2 structure shown in Fig. 1 panel A. Deposited model is in brown and DeepTracer model is in blue. B. Comparison of DeepTracer model with density map. C and D, as in A and B except model-building carried out with the Phenix tool map_to_model and map_to_model structure is in magenta^19^. The unoccupied density in B and D that does not correspond to the brown deposited model in A and C corresponds to an antibody heavy chain that is part of this structure. E. Progress of automated model-building for structures shown in Fig. 1 using AlphaFold prediction iterated with model rebuilding based on a density map. The resolution of the map and the PDB identifier for each structure is listed. The vertical bars show the percentage of C_α_ atoms in the deposited structure that are within 3 Å of any C_α_ atom in the corresponding model. The purple bars represents initial AlphaFold models, superimposed on the deposited structure. The salmon, grey, yellow and red bars respectively, represent the rebuilt model in cycles 1, 2, 3, and 4 of iterative AlphaFold modeling and rebuilding.

The output of the iterative AlphaFold modeling and map-based rebuilding process described above is a predicted AlphaFold model that is already positioned to match the density in a map. The predicted model may still require some adjustment indicated by the density map, and such adjustment can be carried out by automatic refinement^24^ or rebuilding as described above. The resulting refined or rebuilt model is an automatically-generated interpretation of the corresponding part of the density map. Our procedure can therefore also be viewed as a method to automatically interpret a density map, incorporating information from a density map into AlphaFold modeling in the process.

Panel E of Fig. 3 presents results from the same analysis of cryo-EM maps as that shown in Fig. 1, this time from the perspective of automated map interpretation. As in a real case where the structure is not known, each full density map is supplied without any trimming or masking. The sequence of one chain to be interpreted in this map was used to create a standard AlphaFold prediction. That predicted model is automatically oriented to match the map, rebuilt to match the density in the map, and included in the next AlphaFold prediction. After iteration, the last version of the model that was rebuilt to match the map is the output of the procedure. This differs from Fig. 1 in that the final model is now no longer an AlphaFold model, but instead is an AlphaFold model that has been adjusted to match the map. The progress of map interpretation for each of the 25 recent cryo-EM density maps considered in Fig. 1 panel J is shown in Fig. 3 panel E. Some of these structures contain multiple copies of the same chain. In these cases, matching any copy was allowed in this evaluation. Others contained multiple chains with similar sequences (e.g., proteasome structures 7lsx and 7ls5, the antibody heavy and light chains in 7mjs, and the αβγδ histones in 7lv9). In these cases, a match was allowed to whichever chain matched the location the automatic docking had chosen (the correct location was actually picked in all cases except for 7lv9, a structure at a resolution of 4.5 Å). For each structure and density map, panel E shows this percentage of matching C_α_ atoms for the initial AlphaFold model (superimposed on the deposited chain with secondary-structure matching) and the four automatically-docked and iteratively rebuilt models. The structures are arranged based on the resolutions of the corresponding maps, with finer (higher) resolution on the left and coarser (lower) resolution on the right. The SARS Cov-2 spike protein structure^16^ shown in Fig. 1 is labeled as 7mlz in panel E; it can be seen that the automated interpretation of this density map starts with 71% of C_α_ atoms in the deposited model matched by the rebuilt model and improves with each cycle of rebuilding until the next-to-last cycle, where 92% are matched, and no additional improvement is obtained on the final cycle. Others that improve substantially include 7m7b (improving from 59% to 77% matched), 7lx5 (75% to 91%), and 7lci (73% to 95%). In 18 of 25 cases a model matching at least 95% of C_α_ atoms in the deposited structure within 3 Å was obtained; this level of accuracy was present in only 11 of the starting AlphaFold models. Two of the cases (7lv9 and 7msw) yielded very poor models (E). In each of these cases, the initial AlphaFold model was predicted with very low confidence. In the case of 7lv9, the plDDT for only 5 of 97 residues was above the threshold for a “good” prediction of 0.7, for 7msw this portion was 86 of 635 residues. Overall, the accuracy of the 25 chains examined improved from an average of 82% of C_α_ atoms in the deposited model matched of to an average of 91% after iterative modeling and rebuilding.

As the procedures described here are not specific to AlphaFold, to cryo-EM maps or to the Phenix^25^ model rebuilding software used in this work, we expect that the synergy of model prediction and model rebuilding using a density map observed here will be general and that similar results could be obtained using other model prediction and model rebuilding approaches and using other types of density maps such as those obtained in cryo-tomography or crystallography.

## Methods

### Choice of maps and models

The 25 maps and corresponding models shown in Fig. 1 and 3 were chosen in Aug. 2021 in a way that was intended to yield relatively representative recent structures in the PDB. We selected the first protein chain in the 25 most recently-deposited unique cryo-EM structures at the time with resolution of 4.5 Å or better and containing between 100 and 1000 residues. For this purpose, we considered two structures to be duplicates if the first protein chains matched in sequence at a level of 99% identity or greater. We included one pair of similar structures in the 25 structures chosen (7mlz and 7lx5). These differ in residues at the ends of the chain and differ also in that the SARS Cov-2 spike protein (the chain analyzed) is bound to different antibodies in the two structures. The PDB and EMDB accession numbers for these 25 structures are listed in Extended Data Table I.

The choice of structure and map to test model creation using AlphaFold trained without similar sequences in the PDB was made by selecting the (one) structure in CASP-14 that was determined by cryo-EM, classified as a “hard” target in CASP-14^26^, and for which experimental data is available in the EMDB and PDB. This structure was PDB entry 7bgl^20^, EMDB entry 12183. We chose domain 3 of chain a in this structure as AlphaFold performed poorly on this target in CASP-14 (rank of 78) compared to most other targets (rank of 1 for all other 7bgl targets).

### Map and model display

Figures were prepared with ChimeraX^27^.

### Map preparation

For the analyses shown in Fig. 1 and 3, the full maps corresponding to each structure were used. The overall resolution-dependent sharpening or blurring of maps were automatically adjusted using the deposited model with the Phenix tool *local_aniso_sharpen* (without the local sharpening feature but applying the anisotropic correction). For the 7bgl structure in Fig. 2, the map was boxed so as to include the density corresponding to the domain that was analyzed, but was not masked (density corresponding to other chains was therefore present as well).

### Overall procedure for iterative AlphaFold model generation and model rebuilding using a density map

The first cycle of our iterative procedure consists of creating an AlphaFold model using a Google Colab (https://colab.research.google.com/) AlphaFold2 notebook, followed by downloading the resulting model and automatically trimming, docking, and rebuilding the model with the density map and the Phenix tool *dock_and_rebuild*. Subsequent cycles consisted of converting the rebuilt model to mmCIF format^28^, uploading the model to the Colab notebook, generating a new AlphaFold model using the rebuilt model as a template, and rebuilding as in the first cycle. A total of four cycles were carried out. We considered the last AlphaFold model obtained in this procedure to be the AlphaFold model created with information from a density map, and the last rebuilt model to be the overall final model produced by the procedure.

### AlphaFold model generation

We used a slightly modified version of the ColabFold notebook^29^ to create models with AlphaFold. The principal difference from ColabFold is that this notebook can create models for a group of sequences, each with optional uploaded templates. This allowed us to analyze all the structures in Fig. 1 as a group. Another difference is that this notebook allows any combination of use of templates supplied by the user and chosen from the PDB and the optional use of multiple sequence alignments. The notebook is available at https://colab.research.google.com/github/phenix-project/Colabs/blob/main/alphafold2/AlphaFold2.ipynb. In the first cycle of AlphaFold model generation, no templates were used and multiple sequence alignments were included. In subsequent cycles, the rebuilt model from the previous cycle was used as a template. For the examples in Fig. 1, multiple sequence alignments were included in all cycles; for the 7bgl example in Fig. 2, they were included only in the first cycle.

### Automatic model trimming, docking and rebuilding

We used the Phenix^25^ tool *dock_and_rebuild* to orient AlphaFold models in a density map and rebuild them based on the map. This is accomplished in five overall steps: trimming and splitting into domains, docking of individual domains, morphing the full AlphaFold model to match the docked domains, creating rebuilt versions of the model, and assembly of the best parts of the rebuilt versions of the model. All these steps are carried out automatically with the *dock_and_rebuild* tool that in turn uses other Phenix tools to carry out individual steps. Key parameters are noted in the text below; except as noted, default values were used throughout this work.

### Model trimming and splitting into compact domains

AlphaFold models are automatically trimmed and split into domains based on the coordinates of the AlphaFold model and on estimates of confidence (plDDT values^1^) supplied by AlphaFold for each residue in the structure. The Phenix tool *process_predicted_model* is used for this purpose. Residues with plDDT value less than 70 (the threshold for a “good” prediction^1^) are removed and the remaining residues are grouped into “domains” (up to three by default, controlled by the parameter *maximum_domains*) consisting of one or more parts of the chain that contain a sufficient number of residues (10 residues, controlled by the parameter *minimum_domain_length*) and form a compact unit. This grouping can be carried out based on spatial proximity (default), or based on the predicted uncertainties in C_α_ - C_α_ distances. We note that in cycles after the first, a template is supplied that derives in part from the previous AlphaFold model, resulting in systematically higher plDDT values. In this work we have not quantified this effect or adjusted the threshold to account for it.

### Domain docking into density

The compact groups of residues (“domains”) obtained by trimming the AlphaFold model are aligned, one at a time, to the density map. Two approaches are used. The first approach uses secondary structure matching (SSM) to dock the domain onto the map using the Phenix tool *superpose_and_morph* with the setting *ssm_match_to_map=True* (see below for details of this tool). The second approach consists of a direct correlation search between model-based density and the map using the Phenix tool *dock_in_map*^25^. Normally these procedures are carried out sequentially, and if the first yields a match with a map-model correlation (*CC_mask* value using the Phenix tool *map_model_cc)* sufficiently large (typically 0.3, controlled by the parameter *ssm_search_min_cc*) the other is skipped. Based on the hypothesis that the transformations for different domains may often be similar, the transformations for successfully-docked domains are considered as possible transformations for each additional domain. These methods typically yield a set of possible placements of each domain in the map. If symmetry is automatically detected in the map^25^, these placements also include all the possibilities obtained by applying this symmetry to placements found directly.

The final inclusion and placement of each domain is then chosen by maximizing an empirical scoring function. The function includes the fraction of domains that are placed and the map correlation for each placement. It also includes a penalty for placing two domains further apart than can be spanned by the number of residues between those domains, and a penalty function for the number of C_α_ atoms in one domain overlapping with those in another domain within 3 Å (controlled by the parameter *overlap_ca_ca_distance*) The score starts out at zero. If the map correlation for each domain is at least 0.15 *(minimum_docking_cc*) the score is given large positive increases (200 units) for each of the following that occur: (1) lowest map correlation of all docked domains is greater than 0.5 (set with *acceptable_docking_cc*), (2) if #1 occurs and also all placements have similar transformations (i.e., the docking was essentially a rigid-body docking), where two transformations are similar if applying them to a domain gives an rms difference in coordinates equal to the resolution of the map or less, (3) all domains are docked, (4) the fraction of residues that overlap between domains is less than 0.1 (*allowed_fraction_overlapping*), (5) no domains are further apart than can be spanned by the number of residues between those domains. If any domains are further apart than can be spanned by the number of residues between those domains plus twice the resolution plus 15 Å (*maximum_connectivity_deviation*), 200 units are subtracted from the score. The resulting score is then adjusted with the following additions and subtractions: (1) the lowest map correlation of all domains is added, (2) the average map correlation is added, (3) the fraction of transformations that are different from the first is subtracted, (4) the fraction of C_α_ atoms that overlap between domains is subtracted, and (5) the sum of all deviations in distances between domains, normalized to the sum of all allowed distances between domains, is subtracted. This scoring function was not optimized and does not contain weights except as described above.

### Morphing and refining the full AlphaFold model to match the map based on docked domains

Once a set of domains is placed to match a map, the entire AlphaFold model is morphed to superimpose on these domains as much as possible, while smoothly distorting along the chain between domains. We use a shift-field approach to morphing^30^, creating a vector function that varies smoothly in space. The shift (distortion) applied to a particular atom in a model is the value of the shift field at the coordinates of that atom.

The shift field is calculated from a set of (shift coordinate, shift vector) pairs. There is one such pair for each C_α_ atom in a docked domain. The value of the shift coordinate is the position of the corresponding C_α_ atom in the full AlphaFold model. The value of the shift vector is the difference between the coordinate of the C_α_ atom in the docked model and the corresponding C_α_ atom in the full AlphaFold model. The shift field at any point in space is then the weighted average of all the shift vectors, where the weights are the inverse exponential of the normalized squared distance between that point in space and the corresponding shift coordinate, and where the normalization is the square of the shift field distance, which has a typical value of 10 Å (set with the parameter *shift_field_distance* and chosen to be a compromise between maintaining the model geometry with a long shift field distance and matching the docked domains closely with a a short one). The coordinates of a morphed AlphaFold model are then calculated from the initial coordinates and this shift field. This morphing has the property that local distortions occur on a scale of about 10 Å, the shift field distance. The docked, morphed AlphaFold model is adjusted to match the map using the refinement tool *real_space_refine*^24^.

### Creating rebuilt models by replacing uncertain parts of the docked, morphed, refined AlphaFold model

The parts of the docked, morphed, refined AlphaFold model that have either (1) low confidence predictions from AlphaFold (typically residues with plDDT < 0.7 as above), or (2) low correlation with the map, are then identified and used to specify segments of the model that require rebuilding. The threshold defining low map correlation is obtained with the following procedure. Density values in the map at positions of all C_α_ atoms are noted, the values in the lower half are removed, and the mean and standard deviation of remaining (“good”) density values are noted. Low map correlation is defined as more than 3 standard deviations below the mean (where the ratio of 3 is defined by the parameter *cc_sd_ratio*) Before applying these thresholds, the plDDT values and density values for each residue are smoothed by averaging with a window of 10 residues along the chain (defined by *minimum_domain_length*).

Then a series of attempts to improve the fit of each poorly-fitting segment to the map are carried out. These attempts to improve the fit include: (1) iterative resolution refinement, in which the model is iteratively refined, initially at low resolution (6 Å, controlled by the parameter *iterative_refine_start_resolution*), then progressing in 1 Å decrements until the resolution of the map is reached, (2) rebuilding of loops, using the Phenix tool *fit_loops*, (3) retracing loops by finding a path through the density map that connects the ends of the loop with a chain that follows the path with the highest minimum value^19^, (4) a combination of retracing part of the loop with superimposing and splicing that part of the existing refined model that matches the remainder of the loop, (5) iterative morphing, and (6) use of an external model. The combination method addresses the situation where clear density is present in the map for the beginning and end of a loop and the remainder is unclear. In this case, the refined model for the residues that cannot be modeled from density are simply grafted on to the residues that can be modeled, using a shift-field procedure as described above to morph the refined model while superimposing 3 residues on each end. The iterative morphing procedure was similar to one previously used to distort a model to better match the density^31^, but in the current procedure morphing is carried out on 6 residues from each end at a time (specified by *n_window*), then the remainder of the model is superimposed on the 12 morphed residues, the window is shifted by one residue from either end, and the process is repeated until the loop is morphed. In cases where an externally-created model has been supplied to the rebuilding procedure, another attempt to rebuild each loop consisted of selecting a matching segment from the external model, if such a segment with the expected number of residues was present and could be connected to the existing model with deviations at the ends of 3.8 Å or less (defined by the parameter *ca_distance*). Each attempt to rebuild a part of the refined model yields a new candidate segment of the model. All the candidate segments obtained with a particular rebuilding method (e.g., rebuilding loops) are used to replace the corresponding segments in the refined model and the resulting full model is refined based on the density map. This overall process then yields several new full-length versions of the model.

### Assembling the best parts of rebuilt models into a single final model

The rebuilt and refined models are then used as hypotheses for the structure to be built. In the preceding step, boundaries of regions needing or not needing rebuilding were identified. In this step, each model is broken up into the corresponding segments. Then the best version of each segment, chosen based on their map correlation, is used to create a new full model. This model is refined using the density map to produce a single full-length final model.

### Values of parameters

Default values were used for the parameters controlling model rebuilding, with two exceptions. One exception was that for the 7bgl structure^20^ in Fig. 2, model-building was aided by supplying a model created by the Phenix tool *map_to_model^19^* as a source of possible fragments to use in rebuilding the structure. The reason this was necessary was that without these fragments, model rebuilding with the methods described below was incomplete for this structure despite the very good resolution of 2.2 Å, possibly because the AlphaFold model was quite different from the actual structure in some places. The other exception was that in cases where multiple chains with similar sequences (and therefore presumably similar structures) were present in a structure, the secondary-structure-based docking procedure was skipped and only a direct density correlation search was used (with the Phenix tool *dock_in_map*). The rationale for this was that, as might be expected, docking with a correlation search was more effective than a secondary structure search at distinguishing the correct placement from one superimposing on related but different chain.

In the first cycle of rebuilding for each model, the corresponding AlphaFold model was supplied along with the full corresponding density map and the resolution of the structure reported in the PDB. In subsequent cycles, a new AlphaFold model was supplied as well as the rebuilt model from the previous cycle.

### Model superposition and comparisons

Models were superimposed using the Phenix tools *superpose_pdbs*, *superpose_and_morph* and the Coot secondary structure matching tool^32^.

The *superpose_pdbs* tool carries out least-squares superposition of matching C_α_ atoms identified by alignment of the sequences of two models. Note that in cases where the sequences of two models are similar and the models differ largely by rigid-body movement of one domain relative to another, this procedure can lead to a superposition where neither domain superimposes closely.

The *superpose_and_morph* tool carries out secondary structure matching (SSM) to superimpose part or all of one model on another using reduced representations of secondary structure elements and indexing of these elements to speed up comparisons and allowing matches that are nonsequential in a procedure similar to that used in^33^. If the option *ssm_match_to_map* is used, the inputs are a model and a map. In this case the tool *find_helices_strands* is used to find secondary structure elements (SSE’s) in the map and to create a secondary-structure model containing these SSE’s. Then the model to be docked is superimposed on the a secondary-structure model with a modified form of SSM. In this SSM procedure, two secondary structure elements from the map (e.g., a helix and a strand) are paired with two matching elements from the domain to be docked (e.g., a matching helix and strand), thereby defining a transformation between the domain to be docked and the map. As the precise alignment of secondary structure elements from the map and those from the domain to be docked is not known, all possible alignments of the shorter of each pair of elements with the longer element are tested (e.g., residues 1-10 of one helix might be paired with residues 1-10, 2-11, 3-12 and so on from the other). All the C_α_ atoms in each element from the map are then associated with C_α_ atoms in the corresponding element from the domain, and a least-squares superposition is carried out. If these C_α_ atoms match (by default within 5 Å, controlled by the parameter *match_distance_high*), the resulting transformation is applied to all C_α_ atoms in the domain and the map-model correlation of the resulting docked domain is calculated with the Phenix tool *map_model_cc*. If the resulting correlation is above a minimum level (controlled by the parameter *ok_brute_force_cc* with a default value of 0.25), the docked model is adjusted by rigid-body refinement to maximize this correlation.

The Phenix *chain_comparison* tool was used to compare models that were already superimposed. This tool counts the number of C_α_ atoms in a target model that are matched within 3 Å by any C_α_ atom in the matching model. Allowing any C_α_ atom in the matching model to superimpose effectively ignores the connectivity of the chains, but it is useful for evaluating whether a C_α_ atom is placed in a position where some C_α_ atom belongs. The distance of 3 Å is the default value and is useful for ranking pairs of models that have more than about 30% of C_α_ atoms matching. It is less useful for ranking pairs with lower similarity because two overlapping structures that are completely unrelated will often have 20-30% of C_α_ atoms matching within 3 Å.

### Map correlations

We used the Phenix tool *map_model_cc* to calculate correlations between experimental density maps and model-based density maps for PDB entry 7mlz and resulting AlphaFold and rebuilt models. The overall orientation and positions of AlphaFold models are arbitrary and the values in the atomic displacement parameter field (B-values) are plDDT values. We superimposed these models on the corresponding deposited structure before calculation of map correlations, keeping all coordinates fixed at the values obtained by direct superposition. To make a fair comparison with rebuilt and deposited models, we refined the atomic displacement parameters for all the models to match the map before calculation of map correlations. For the 7mlz example shown in Fig. 1, the refinement of B-values increased all the map correlation values. Map correlation values for the deposited model, initial superposed AlphaFold model (with B-values representing plDDT), initial rebuilt model, and final AlphaFold model were 0.64, 0.26, 0.47, and 0.44. After B-value refinement these were 0.70, 0.41, 0.58, and 0.57.

## Acknowledgments

The authors appreciate funding from the Department of Energy National Nuclear Security Administration (grant DE-AC52-06NA25396 to PDA), from Lawrence Berkeley National Laboratory (grant DE-AC02-05CH11231 to PDA), from the Phenix Industrial Consortium (to PDA), and from the National Institutes of Health (grant GM063210 to PDA, RJR, TCT).

## Author contributions

The overall concepts in the paper were developed and supervision was carried out by TCT, PDA, RJR, and JSR. TCT wrote the initial draft and carried out the analyses. BKP, PVA, CJS, TIC and CM developed tools that were essential to the work. All authors contributed ideas to the work and assisted in editing of the manuscript.

Authors declare that they have no competing interests

## Additional Information

### Data Availability and Code Availability

All maps and models used in this work were obtained from the PDB and the EMDB. All code for the Phenix version of the AlphaFold2 Colab is freely available from github at git@github.com:phenix-project/Colabs.git. All code for Phenix is available at phenix-online.org. The spreadsheets and ChimeraX sessions used to generate the figures in this paper, along with the maps and models created for Figs, 1, 2 and 3 are available at: https://phenix-online.org/phenix_data.

Reprints and permissions information is available at www.nature.com/reprints.

Supplementary Information is available for this paper

## Extended Data

### Structures and maps used

**Extended Data Table I.** List of structures and maps used in Figs. 1 and 3. The most similar entries at time of deposition were obtained using the RCSB PDB “Find similar proteins by 3D structure tool” and choosing the highest-scoring entry that was deposited earlier than the target structure. The percentage of residues matching and rmsd were obtained using the Phenix superpose_pdbs tool using only C_α_ atoms from the target and noting the number of residues in the target, the number of superposed residues, and the final rmsd of matching C_α_ atoms.

| **PDB ENTRY** | **EMDB ENTRY** | **RESOLUTION (Å)** | **CHAIN** | **MOST SIMILAR IN PDB (at time of deposition)** | **Percent matching** | **RMSD (Å)** |
| --- | --- | --- | --- | --- | --- | --- |
| 7mby | 23750 | 2.44 | B | 6pt0 | 2 | 19.5 |
| 7me0 | 23786 | 2.48 | A | 7k0r | 98 | 0.4 |
| 7ev9 | 31325 | 2.6 | A | 1yew | 100 | 0.5 |
| 7ls5 | 23502 | 2.74 | A | 3gpt | 99 | 0.4 |
| 7eda | 31062 | 2.78 | A | 5xnl | 4 | 21.6 |
| 7lci | 23274 | 2.9 | R | 6lmk | 7 | 4.1 |
| 7lvr | 23541 | 2.9 | A | 6b0i | 100 | 1.0 |
| 7c2k | 30275 | 2.93 | A | 7btf | 98 | 1.0 |
| 7m7b | 23709 | 2.95 | A | 7m3i | 98 | 2.9 |
| 7n8i | 24237 | 3 | L | 6iw2 | 3 | 2.6 |
| 7mjs | 23883 | 3.03 | H | 6ukj | 2 | 7.0 |
| 7l1k | 23110 | 3.16 | A | 4kvo | 2 | 14.6 |
| 7l6u | 23208 | 3.3 | A | 5hej | 94 | 1.3 |
| 7ku7 | 23035 | 3.4 | A | 5ejk | 91 | 1.7 |
| 7kzz | 23093 | 3.42 | B | 5vrf | 98 | 1.4 |
| 7lx5 | 23566 | 3.44 | B | 7klh | 2 | 5.7 |
| 7brm | 30160 | 3.6 | A | 4uv3 | 92 | 0.7 |
| 7lsx | 23508 | 3.61 | A | 6fvw | 8 | 23.7 |
| 7lc6 | 23269 | 3.7 | A | 6hrb | 1 | 10.2 |
| 7mlz | 23914 | 3.71 | A | 7lrs | 2 | 15.5 |
| 7msw | 23970 | 3.76 | A | 6ce1 | 2 | 17.0 |
| 7rb9 | 24400 | 3.76 | B | 6upv | 98 | 0.4 |
| 7bxt | 30237 | 4.2 | A | 6mup | 22 | 0.8 |
| 7m9c | 23723 | 4.2 | A | 3udb | 1 | 6.0 |
| 7lv9 | 23530 | 4.5 | B | 5x0y | 5 | 16.4 |

**Extended Data Figure 1.**
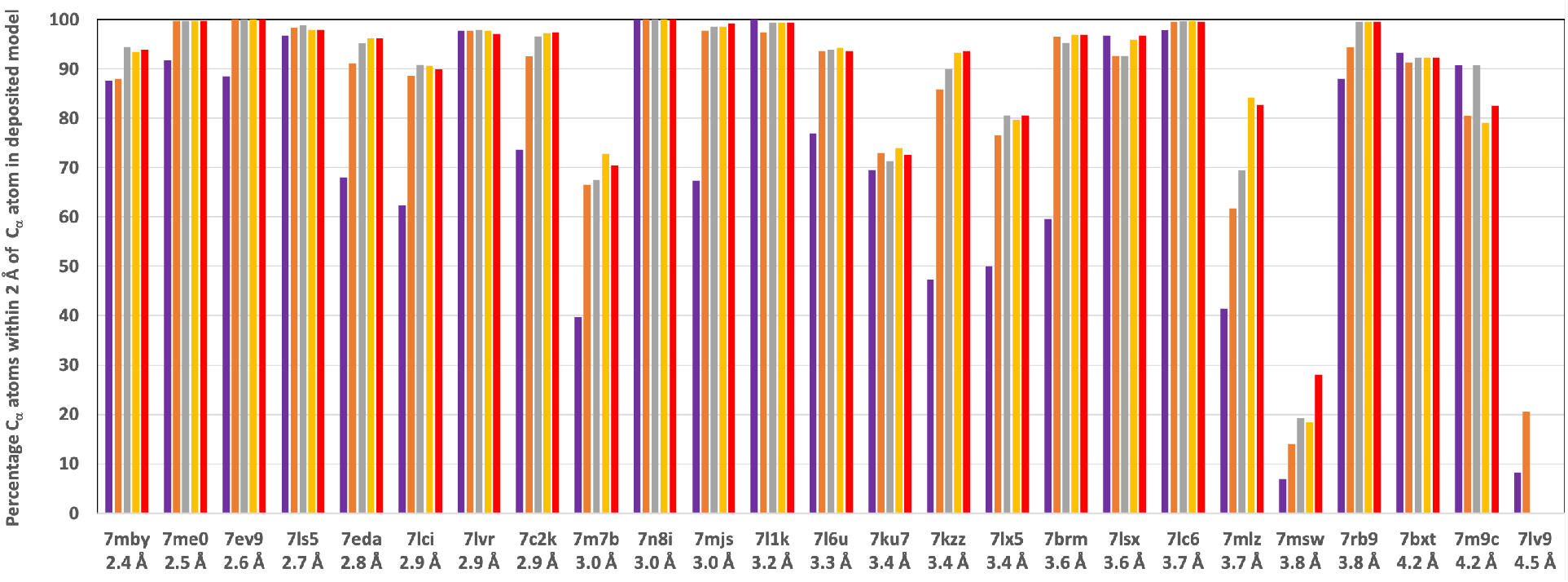
Analysis of automatic map interpretation in Fig. 3 panel E using 2 Å cutoff instead of 3 Å. Figure as in Fig. 3 panel E, except the value of the keyword max_dist was set to 2 Å instead of 3 Å. This then reports the percentage of C_α_ atoms in the deposited structure matched within 2 Å by a C_α_ atom in the corresponding final rebuilt model.

## Supplementary Information

### Prediction of 7bgl flagellar basal body structure as a homo-multimer

The flagellar basal body structure is a symmetric 26-mer (C26), while the AlphaFold prediction in CASP-14 used a monomer, and we also used the sequence of a monomer in our analysis. It seemed likely that some of the sequence covariation present in the multiple sequence alignment would be due to inter-subunit contacts, and that if we supplied a sequence corresponding to a homo-oligomer, AlphaFold might be able to use this inter-subunit contact information to create a more accurate model of each individual chain. We carried out a prediction with a 26-mer of the 7bgl sequence; this model had highly overlapping copies of the chain and was not considered further. We also carried out predictions with a trimer and a dodecamer of the 7bgl sequence and compared one chain from each with the deposited model. The conformation of each chain in predictions from the trimer were similar to that of the predicted monomer in Fig. 2 panel A, and had an rmsd from the deposited model of 3.8 Å. Three of five dodecamer predictions had a conformation for each single chain that was more like the deposited model, with an rmsd of superposed C_α_ atoms for the top-scoring model of 3.0 Å. For comparison, the iterative AlphaFold model in Fig. 2B has an rmsd of 0.8 Å.

### Iterative model improvement requires both iterative rebuilding and iterative modeling

We carried out tests to check whether the improvement obtained between Fig. 1 panels A and C actually requires both the iteration of prediction and the iteration of model rebuilding. To test whether model rebuilding is necessary, we carried out iterative cycles of AlphaFold prediction for the SARS Cov-2 spike protein^1^ example in Fig. 1 panel A using the AlphaFold model from each cycle directly as a template in the next. The resulting prediction after four cycles was very similar to the original AlphaFold model, with an rmsd of matching C_α_ atoms in the model of just 0.5 Å after superposition, and differed very substantially from the model obtained with iterative prediction and rebuilding (corresponding rmsd of 3.2 Å), indicating that without model rebuilding, iteration of the procedure has little effect.

To test whether iteration of prediction is necessary, we carried out iterative cycles of rebuilding of the same structure, starting with the rebuilt model in purple in Fig. 1 panel B. This starting model had an rmsd of matching C_α_ atoms in the deposited model of 5.7 Å, while the model in Fig. 1 panel C had a corresponding rmsd of 2.4 Å. Carrying out iterative cycles of rebuilding without further prediction yielded a model with a much greater rmsd to the deposited model of 4.4 Å, showing that without iteration of the prediction part of the process, the procedure is not effective.

### Situations where non-default choices of parameters may be useful in modeling

The principal options that are available in our procedure for iterative AlphaFold prediction and rebuilding are (1) including an external model as a source of hypotheses during model rebuilding, and (2) omitting multiple sequence alignments in cycles of AlphaFold after the first.

The use of an external model may be useful in cases where the density map is clear but the automatically docked and trimmed AlphaFold predicted model matches the map poorly enough that the automatic rebuilding process fails. This can happen for example if the AlphaFold model is placed incorrectly in the map or if the residues remaining at the ends of trimmed fragments agree so poorly with the density map that the rebuilding process is not able to identify connections between them. An indication for using this approach is that the density map appears to show clear density for a protein but neither the rebuilt model nor the final AlphaFold model matches that density. An external model from an automatic map interpretation procedure such as Phenix *map_to_model*^2^ or DeepTracer^3^ could be used as well as a manually built model.

Omitting multiple sequence alignments after the first AlphaFold cycle may be useful in cases where the sequence alignment causes AlphaFold to create a model that is inconsistent with the density map. The residue covariation deduced from multiple sequence alignments are presumed to be a principal source of information about residue-residue distances for AlphaFold ^4^ but at the same time multiple sequence alignments may have significant uncertainties^5^. If a multiple sequence alignment is inconsistent with the actual structure of the protein being modeled, inclusion of this alignment could make modeling work poorly. In AlphaFold modeling the contribution from a multiple sequence alignment is often very substantial^4^, but in cycles after the first our procedure supplies a template and use of the multiple sequence alignment is less critical and can be omitted. An indication that this option might be useful would be that the rebuilt model produced by the *dock_and_rebuild* procedure matches the map but the resulting AlphaFold model does not.

## Notes

https://phenix-online.org/phenix_data/terwilliger/alphafold_with_density_2022/

### Competing Interest Statement

The authors have declared no competing interest.

### Summary of Updates

The paper is updated to list the most similar entry in the PDB at time of deposition of the 25 structures analyzed and to include a version of Fig. 3E using a cutoff of 2 A as suggested in an on-line review of the manuscript. Additionally another example of major improvement in a structure is shown in Fig. 1 panels G-I.

